# rDNAcaller: a fast and robust pipeline to call ribosomal DNA variants

**DOI:** 10.1101/2025.05.13.653643

**Authors:** Jose Miguel Ramirez, Winona Oliveros, Raquel García-Pérez, Alba Jimenez-Lupiañez, Aisha Shah, Pablo Perez-Cano, Fairlie Reese, Miguel Vazquez, Marta Melé

## Abstract

Ribosomal DNA (rDNA) genes are essential components of the ribosome, organized as tandem repeats on the mammalian genomes. Extensive genetic and copy number variation in rDNA has been reported both within and between individuals, contributing to phenotypic diversity. However, previous rDNA variant calling strategies have relied on methods designed for diploid regions and have not been systematically benchmarked. With the recent availability of a telomere-to-telomere (T2T) genome assembly, rDNA regions have been fully assembled for the first time, enabling benchmarking and optimization of rDNA variant calling strategies. We developed a customized simulator that replicates real intra- and inter-individual rDNA variation based on the T2T assembly to benchmark the performance of commonly used variant callers, including GATK, Mutect2, and Lofreq. Additionally, we optimized the preprocessing and mapping steps to remove pseudogenes and significantly improve accuracy. Based on these optimizations, we introduce *rDNAcaller*, a novel pipeline for accurate rDNA variant detection using short-read whole-genome sequencing. *rDNAcaller* integrates optimized preprocessing with the top-performing variant caller and achieves 94% precision on experimental data from the T2T cell line. Applying our pipeline to data from the 1000 Genomes Project, we identify 5,607 novel rDNA variant positions across human populations, with African individuals showing the highest number of variants. Overall, *rDNAcaller* is a robust and versatile tool for analyzing rDNA variation, addressing the limitations of existing methods in handling high ploidies. By enabling accurate detection of rDNA variants, it facilitates deeper exploration of rDNA’s role in phenotypic diversity, supporting future genomic studies and broadening our understanding of rDNA biology in health and disease.

**Graphical abstract:** 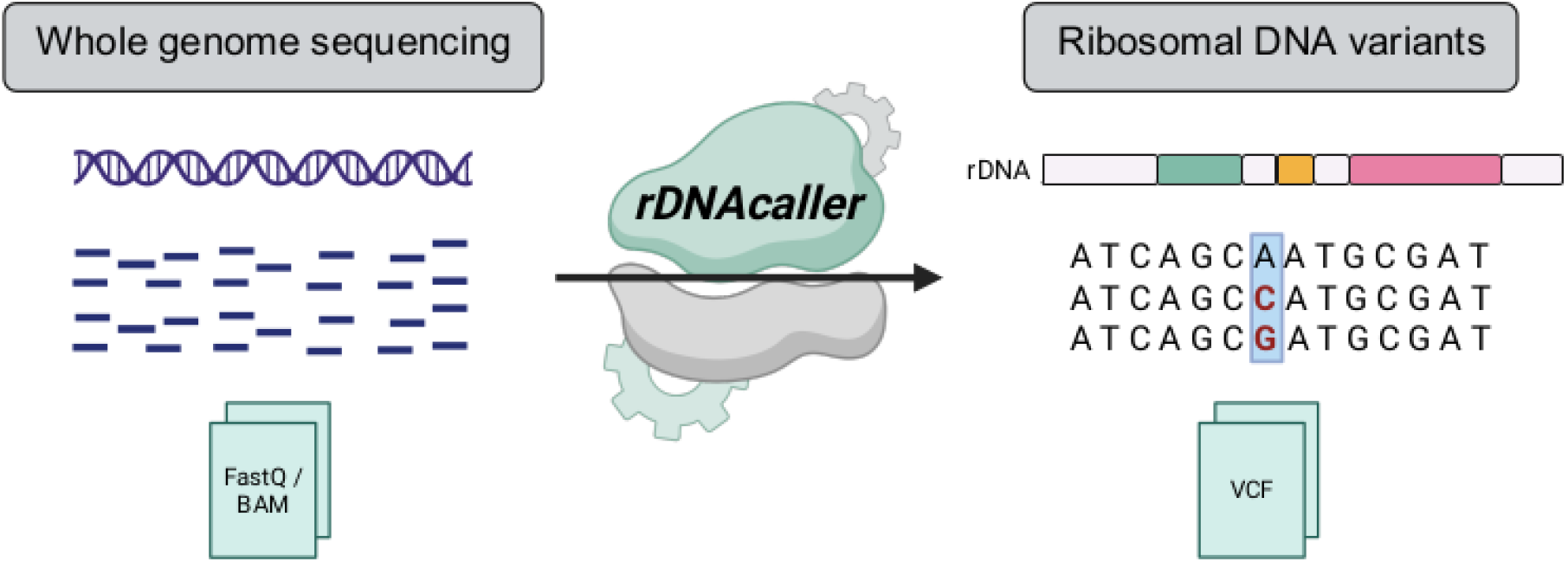

## Introduction

The ribosome is the cellular machinery responsible for protein synthesis. It is composed of ribosomal proteins and ribosomal RNAs (rRNAs). The 5S rRNA is encoded in chromosome 1. The 18S, 5.8S and 28S rRNAs are encoded together by the 45S ribosomal DNA (rDNA), which is organized in tandem repeats on the short arms of human acrocentric chromosomes (13, 14, 15, 21, and 22). (Fig. 1A) (1, 2). The total number of these rDNA repeats, known as rDNA copy number, varies widely among individuals, ranging from 100 to 600 copies (3, 4).

**Figure 1.**
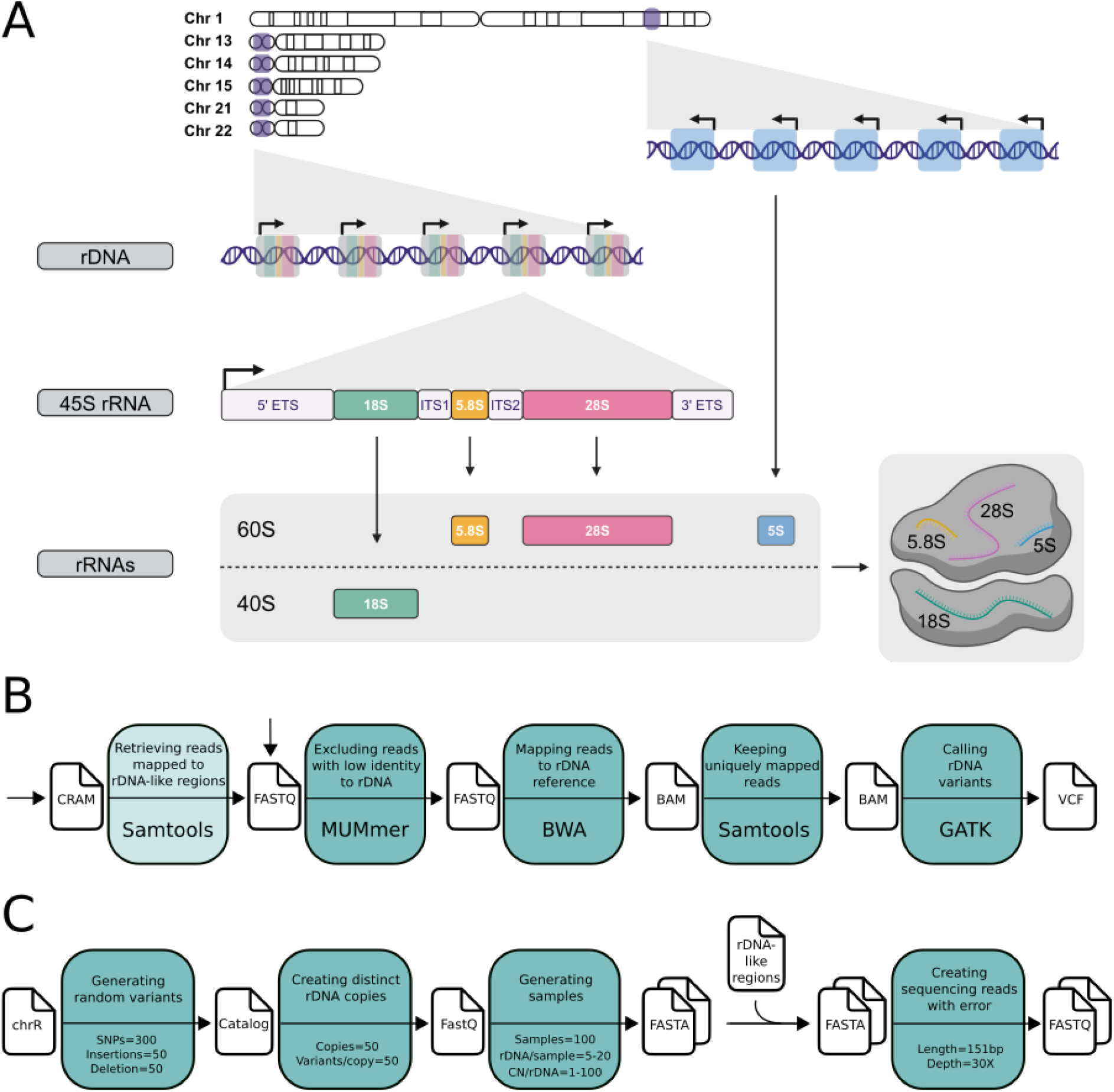
rDNA organization and pipeline diagrams. **A**. Overview of the ribosomal DNA chromosomal organization. rDNA genes are located in the short arms of the acrocentric chromosomes in tandem repeats. The 45S rRNA operon is transcribed as a single transcript and then 18S, 5.8S and 28S are post-transcriptionally processed to be incorporated into the ribosome as separate transcripts. The external and internal transcribed spacers (ETS/ITS) are not incorporated in the ribosome. The 5S is encoded separately in chromosome 1. **B**. Processing steps of the *rDNAcaller* pipeline. Lighter color boxes indicate an optional step which depends on the starting input file (either CRAM or FASTQ) **C**. rDNA read simulator pipeline.

Notably, the rDNA copies are not identical, resulting in substantial inter- and intra-individual genetic variation (5, 6). Different rDNA variants have been shown to have tissue-specific expression patterns (7, 8) and to be associated with specific phenotypes, such as body size (9). However, previous efforts to call rDNA variants have lacked comprehensive benchmarking (6–8, 10) and therefore focus mostly on common variants. This means that the effect of population on rDNA variation and the association between rDNA copy number and rDNA genetic variation remain underexplored (11). This limitation is primarily due to the absence of datasets with known rDNA sequences that could be used as gold standards, since genome assemblies have historically excluded highly repetitive regions such as rDNA. The advent of long-read sequencing technologies has recently enabled the assembly of the first complete telomere-to-telomere human genome (5). This assembly includes the rDNA regions, allowing for the first time the benchmark of a pipeline designed to call rDNA variants.

Here, we present *rDNAcaller*, a pipeline designed for accurate rDNA variant calling from short-read sequencing data. We first create a custom simulator to generate rDNA reads from samples with diverse rDNA copy numbers and genetic variants. We use this simulated dataset to benchmark three variant callers and optimize our pipeline. *rDNAcaller* integrates fast preprocessing and mapping strategies that exclude rDNA pseudogenes and employs the top-performing rDNA variant caller, achieving an F1 score of 96%. Additionally, we evaluate the pipeline’s performance using WGS from the same cell line of the T2T assembly, for which the rDNA sequences are assembled and achieve 94% accuracy. Finally, we apply *rDNAcaller* on the 1000 Genomes Project dataset, identifying thousands of novel rDNA variants across human populations.

## Methods

### Pre-processing and mapping in *rDNAcaller*

The first step of the *rDNAcaller* pipeline is to extract candidate rDNA reads from the FASTQ files of whole-genome sequencing (WGS) data using the function *nucmer* from MUMmer (12) (Fig. 1B). This function parses a FASTQ file and selects the reads that match a reference sequence contiguously for a minimum nucleotide length. The *rDNAcaller* pipeline selects only paired-end reads that both match at least 30 consecutive nucleotides to the rDNA reference sequence using the parameters “*-maxmatch -l 30*”. The rDNA reference sequence, named chromosome R (chrR), has a total length of ~45 kb, including the 13.4 kb transcribed region, the 45S rRNA (13). This MUMmer step speeds up the mapping and variant calling while removing reads from rDNA pseudogenes (see Results).

We then map the selected candidate rDNA reads to a modified version of the CHM13-T2T genome (5), specifically designed to analyze rDNA (13). This is a publicly available modified CHM13-T2T genome reference that contains all rDNA-like regions masked, and a single rDNA reference sequence appended as a separate chromosome, the chrR (13). We map the candidate rDNA reads using *bwa mem* (Li et al. 2013) and we use *samtools* (14) to keep the uniquely mapped and properly paired reads to the chromosome R rDNA reference sequence.

### rDNA sequence simulator

To benchmark available variant callers and include the top-performing one in *rDNAcaller*, we created an rDNA sequence simulator capable of closely replicating true rDNA variation (Fig. 1C). In each run, the simulator first creates a catalog of random variants on the chrR reference sequence (13). The number of variants, along with all other parameters, can be specified by the user. Specifically, we generate 300 single nucleotide variants (SNVs), 50 insertions and 50 deletions of size ranging from 2 to 7 base pairs. Next, the simulator creates 50 distinct rDNA sequences, each containing 50 random variants from the catalog. From this catalog, it then generates 100 samples, each containing between 5 to 20 distinct rDNA sequences, with copy numbers ranging between 1 to 100 copies per distinct sequence. The total copy number per sample is constrained between 100 and 600 copies. The output of this step is a FASTA file for each sample containing all generated rDNA sequences. In addition, the simulator adds all rDNA-like regions of the hg38 genome reference, which include pseudogenes described in (13) to each generated FASTA file. This step ensures that both rDNA copies and rDNA pseudogenes are simulated, reflecting real genomic conditions where the presence of pseudogenes may lead to increased false positives. Finally, we use NEAT (15) to simulate Illumina next-generation sequencing reads from the generated FASTA files, incorporating sequencing errors. Specifically, we simulate 151 bp paired-end reads at 30X coverage.

We ran 50 simulations with the parameters described above, which were selected to reflect rDNA variation observed in experimental data (3, 4).

### Benchmarking variant callers using simulated data

None of the publicly available variant callers are optimized for calling rDNA variants. This is due to rDNA having effectively high and individually-variably copy numbers (high ploidies).

Using our simulations, we benchmarked three publicly available and widely-used variant callers: *HaplotypeCaller, Mutect2* and *LoFreq. HaplotypeCaller* and *Mutect2* are part of the GATK workflow (16) but *HaplotypeCaller* is optimized for germline mutations and allows the analysis of different ploidies, whereas *Mutect2* is optimized for somatic mutations. *LoFreq* is also a variant caller optimized for somatic mutations (17) and has been the most widely-used in previous studies to analyze rDNA variation (7, 10).

We run *HaplotypeCaller* from GATK (16) with different --sample-ploidy (2, 5, 10, 20, 30, 40) and hard filtering of quality > 1000 on our simulated data. The --sample-ploidy parameter in GATK refers to the number of chromosome copies expected in an individual sample, which is usually 2 in diploid loci but in rDNA, the true ploidy is highly variable between samples.

In addition, while changing the parameter --sample-ploidy, we also change --max-genotype-count, an internal parameter, accordingly. The default internal parameter in GATK for --max-alternate-alleles is 6 and for --max-genotype-count, 1024. The number of --max-genotype-count needs to be updated with changing ploidies according to the following formula:

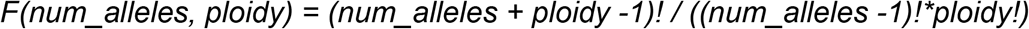

If --max-genotype-count is not updated when increasing --sample-ploidy, multiallelic positions variants will not be evaluated. Hence, we used as --max-genotype-count the result of the formula to keep constant num_alleles=6. For example, for --sample-ploidy 20, we use --max-genotype-count 53130.

Next, we run joint genotyping with GATK, which leverages population-wide information when multiple samples are available. After executing *HaplotypeCaller*, instead of applying a hard filter, we use *GenomicsDBImport* and *GenotypeGVCFs* with the recommended hard filters according to the GATK best practices: for SNPs we use *“QD < 2.0, QUAL < 30.0, SOR > 3.0, FS > 60.0, MQ < 40.0, MQRankSum < −12.5, ReadPosRankSum < −8.0”*, and for INDELs, *“QD < 2.0, QUAL < 30.0, FS > 200.0, ReadPosRankSum < −20.0”*.

Finally, we used two methods optimized for somatic mutations. We run *Mutect2*, a variant caller within GATK (16), with parameter “*--mitochondria-mode*”, which is optimized for variant calling in mitochondrial DNA, a scenario with ploidies similar to rDNA. We also used LoFreq (17) with default parameters.

To compare the different variant callers we computed the following statistics:

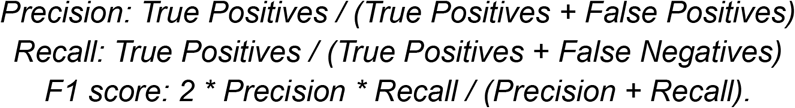

We included the novel preprocessing steps and the best-performing variant caller, *HaplotypeCaller* with ploidy 20, inside a pipeline named *rDNAcaller*, which is publicly available on github: https://github.com/Mele-Lab/rDNAcaller.

Since variant calling with short-read sequencing has been associated with low precision in highly-repetitive regions (18), we analyzed the performance of *HaplotypeCaller* in rDNA repetitive regions and compared it to the rest. To identify repetitive regions, we used the union of Repeatmasker (19) with parameters “*-species human -norna”*, Tandem Repeat Finder (20) with parameters “*2 7 7 80 10*

*50 500 -f -d -m*”, and the homopolymers longer than 3 bases identified with *homopolymerFinder* from sarlacc (21).

### rDNA variant extraction in the human CHM13-T2T assembly

The CHM13-T2T genome assembly contains all rDNA copies present in the CHM13 cell line. To evaluate the performance of *rDNAcaller* in experimental data from the CHM13 cell line, we needed to extract the rDNA variants from the genome assembly. We obtained the genomic coordinates of the rDNA copies from the genome annotation file and extracted them using the function *getfasta* from *bedtools (22)*. To obtain the number of unique rDNA copies we performed a multiple sequence alignment using *clustalo (23)* with parameters “*--full --percent-id --iterations 50*”. We kept the copies without other identical copies based on the distance measures obtained in the multiple sequence alignment. To get the list of variant positions indexed according to chromosome R, we performed a multiple sequence alignment for the 24 unique rDNA copies and the 13 kb of the 45S in chromosome R with the same parameters as before. We extracted the rDNA variants (.vcf file format) from the multiple sequence alignment (.aln file format) using *msa2vcf* from https://github.com/pinbo/msa2snp. Since the indels are reported differently by *msa2vcf* and GATK, we only kept SNPs when comparing the performance of different preprocessing steps on samples from the CHM13 cell line.

### rDNA variant calling in the 1000G

We used *rDNAcaller* in the 1000G project dataset (24). We downloaded the CRAM files from the 2,504 individuals included in Phase 3 from their portal. We excluded 10 individuals with < 300 rDNA reads mapped to chrR.

When alignment files (cram/bam/sam) with reads mapped to *hg19* (*GRCh37*), *hg38* (*GRCh38*) or *hs1* (*T2T-CHM13*) are available, we can directly extract into a FASTQ file the reads mapped to regions of the reference genomes annotated as rDNA-like regions. These are all the regions of the reference genome where an rDNA read could map: 361 kb, 881 kb, and 12 mb for hg19, hg38, and hs1, respectively (13). This is much faster than converting the full cram/bam/sam files into FASTQ.

### rDNA copy number estimation

We assessed rDNA copy number (CN) in the individuals from the 1000G project (25). Briefly, we sorted, indexed and filtered the alignment files (BAM) previously generated by applying our *rDNAcaller*, to retain only reads aligned to the 18S reference using samtools version 1.10 (26). Note that CN is usually computed from reads aligned to 18S instead of 28S, as this second contains higher GC content. We then divided the number of reads mapped to the 18S by the sum of reads assigned to numbered chromosomes as reported by samtools idxstats to obtain the 18S Ratio. Finally, we calculated CN estimates by the following formula:

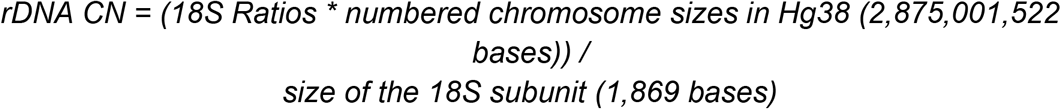

### Nucleotide diversity score

Nucleotide diversity corresponds to the average number of nucleotide differences per position between every sequence pair in a population. This would correspond to the probability of detecting two different nucleotides at the same position from two different sequences. We computed nucleotide diversity scores (π) accounting for polyploidy and multiallelic positions. We computed the mean allele frequencies per variant position across donors and computed all pairwise combinations of allele products, which is equivalent to:

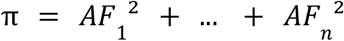

where n is the number of alleles per position. Finally, we computed the average π per region.

### Association between rDNA variants and population

To test the association between number of rDNA variant alleles and population, we fitted a generalized linear model with a quasi-Poisson error distribution to account for overdispersion in the response variable, using the *glm* function from R. We corrected for rDNA CN as a continuous variable and included population as a categorical variable:

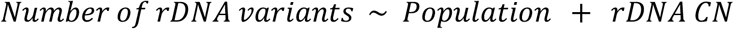

To then test the association between nucleotide diversity score and population, we fitted a linear model using the same covariates as before with the function *lm* from R:

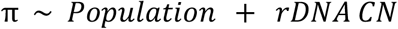

To visualize the results, we computed the residuals with the function *residuals* from R using the same two models without the variable Population.

## Results

### *HaplotypeCaller* with high ploidy is the best-performing rDNA variant caller

To build a pipeline for rDNA variant calling, we benchmarked different variant caller workflows using data generated with our rDNA simulator (Fig. 1C). None of the standard variant caller workflows are optimized to call variants in genes with very high copy numbers, let alone to do so in samples with variable copy numbers as is the case for rDNA genes. We benchmarked three variant callers: GATK’s *HaplotypeCaller*, a method optimized for germline variants that has a ploidy parameter; *Mutect2*, a method optimized for mitochondrial variants; and *LoFreq*, which is optimized for somatic mutations. We also evaluated if running *Joint-Genotyping* after running *HaplotypeCaller* improves performance, as GATK recommends to leverage population-wide information from multiple samples (16).

We benchmarked all three variant callers against 50 different sets of simulations. *HaplotypeCaller* has the highest precision (Fig. 2A) whereas *LoFreq* and *Mutect2* have the highest recall (Fig. 2B). Based on F1 score, *HaplotypeCaller* is the best-performing variant caller (Fig. 2C). Although GATK recommends *Joint-Genotyping* when having multiple samples, in this context, it achieves the lowest performance of all rDNA variant calling approaches. Increasing the ploidy parameter in *HaplotypeCaller* reaches a plateau at ploidy 20 (mean F1 score = 0.96). Thus, increasing the ploidy parameter further only increases computing power with no accuracy benefit. Therefore, we considered GATK’s *HaplotypeCaller* with ploidy 20 as the best-performing rDNA variant caller. Notably, this approach outperforms previously used methods. *Joint-Genotyping* with ploidy 2, the method used by (6) has a mean F1 score of 0.74, and *Lofreq*, the method used by (7), (10) and (9) has a mean F1 score of 0.83 (Fig. 2C).

**Figure 2.**
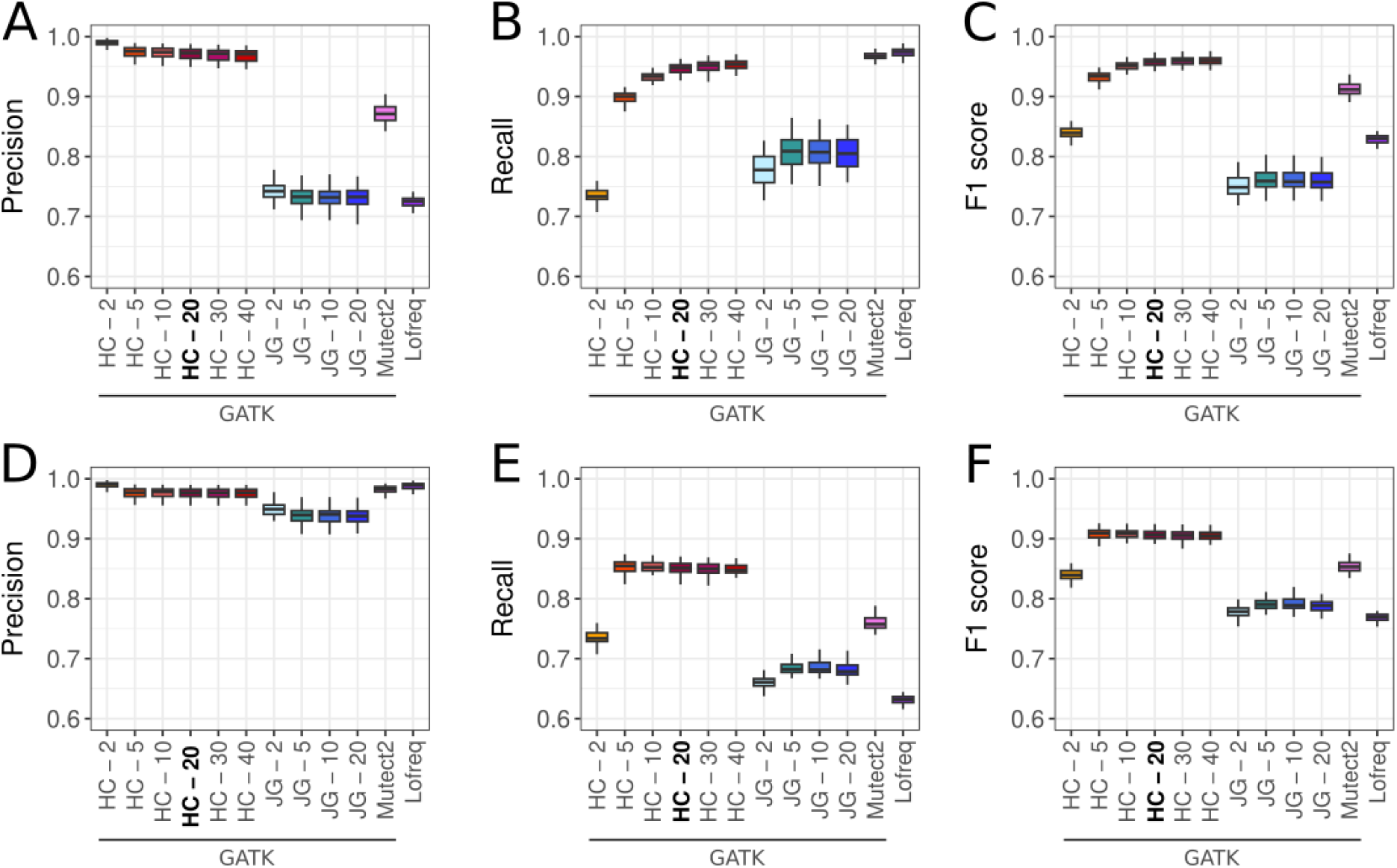
Benchmark of variant callers with rDNA simulated data. **A**. Precision **B**. Recall and **C**. F1 score of the different variant caller and parameters benchmarked using 50 simulations. **D**. Precision **E**. Recall and **F**. F1 score evaluated only using variants with allele frequency > 0.05. **HC** = GATK’s HaplotypeCaller with changing ploidy parameter. **JG** = GATK’s joint genotyping. JG - 2 was the method used by (6) and Lofreq by (7), (10) and (9).

A common practice in previous studies without benchmarking was to filter variants based on allele frequency (9, 10). When filtering out variants with allele frequency lower than 0.05, the precision is very high with all methods (Fig. 2D) to the detriment of the recall (Fig. 2E). Nevertheless, *HaplotypeCaller* remains the top-performing variant caller (Fig. 2F).

Variant calling using short-read sequencing has been associated with low precision in highly-repetitive regions (18). Since rDNA contains highly-repetitive regions, we evaluated the performance of *HaplotypeCaller* in the subset of positions that fall in repetitive regions (Fig. 3A). The F1 score in repetitive regions (mean F1 score = 0.93, Fig. 3B) is slightly lower than when considering all positions (mean F1 score = 0.96, Fig. 3C). However, this decrease is mostly driven by a decrease in recall, and not a decrease in precision (Fig. 3B-C). This suggests that while the number of false negatives might be slightly higher in the rDNA repetitive regions, the number of false positives is not higher. Thus, we can confidently trust the variants called by *HaplotypeCaller* across all the rDNA positions.

**Figure 3.**
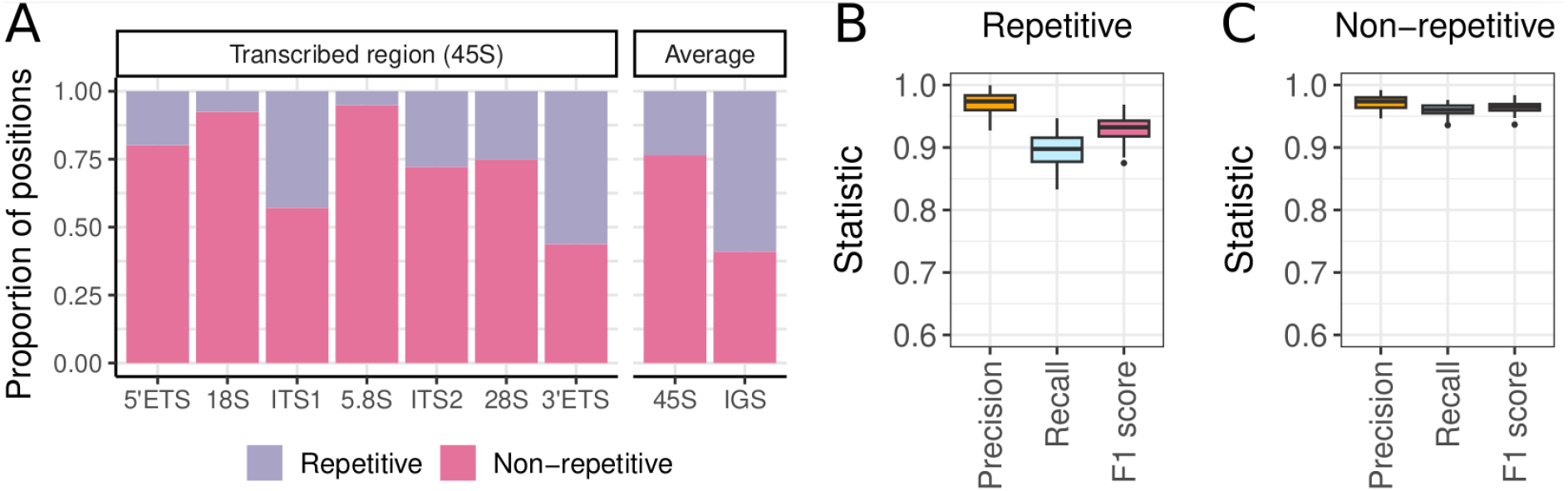
Performance of HaplotypeCaller in repetitive regions. **A**. Proportion of rDNA nucleotide positions that are located in repetitive versus non-repetitive regions. **B**. Precision, recall and F1 score of *HaplotypeCaller* with ploidy 20 in repetitive regions, or **C**, non-repetitive regions.

### *rDNAcaller* is the fastest pipeline

We named our pipeline *rDNAcaller*, which includes selecting candidate rDNA reads, mapping to a customized reference genome, and performing rDNA variant calling with *HaplotypeCaller* ploidy 20 (Fig. 1B). We tested the performance of *rDNAcaller* on experimental data. Specifically, we used short-read WGS from the CHM13 human cell line for which the rDNA copies are present in the T2T genome assembly (5). This cell line contains 219 rDNA copies, from which there are 24 unique rDNA sequences. These unique sequences are extremely similar to one another (Fig. 4).

**Figure 4.**
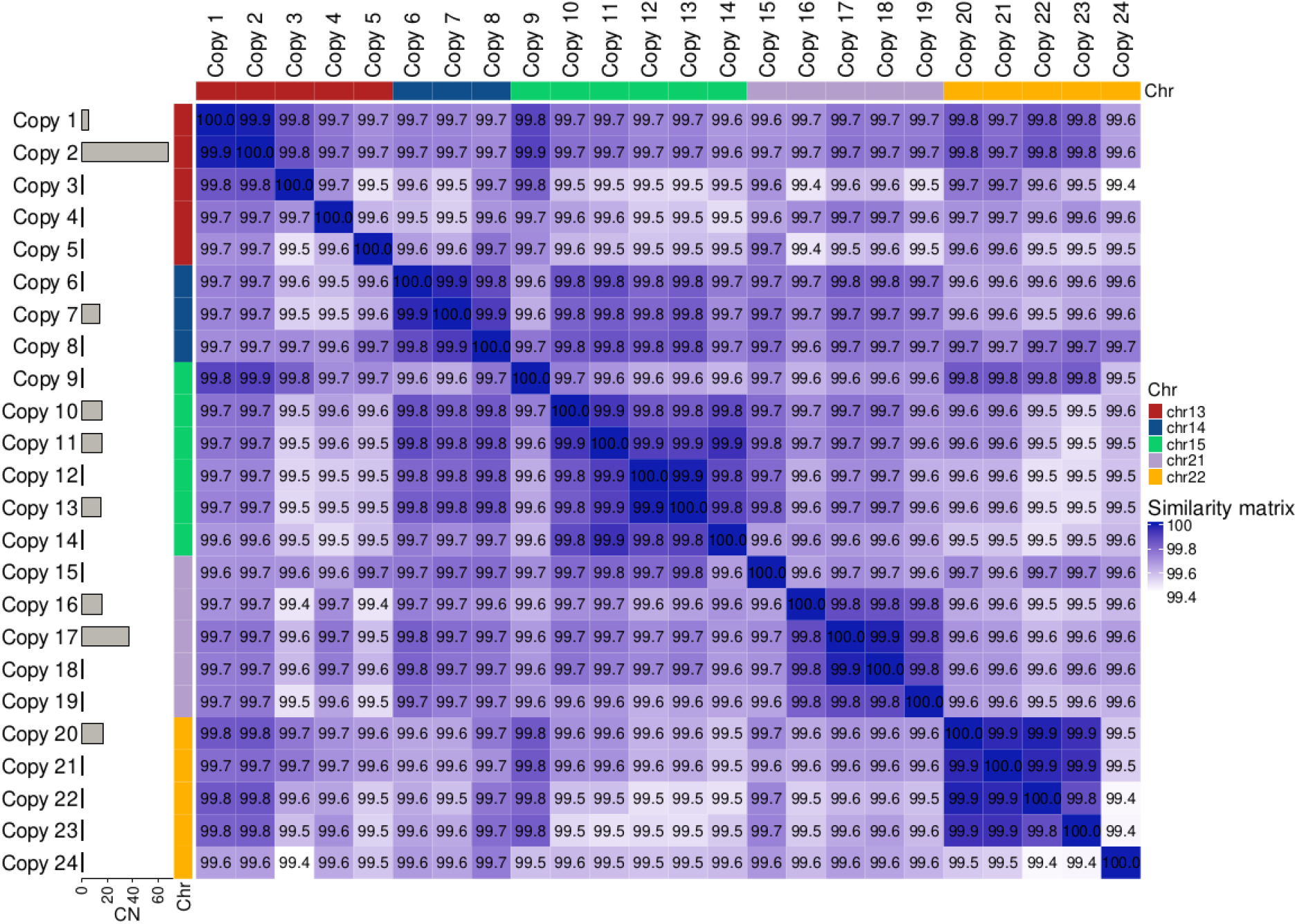
Human T2T unique rDNA copies. **A**. Percentage of similarity between the 24 unique rDNA copies in the CHM13 cell line.

We evaluated the speed and accuracy of our preprocessing steps relative to previously used approaches in the literature. We specifically evaluated two tasks: (1) the inclusion of the MUMmer step, which retrieves candidate rDNA reads with a 30 nucleotide exact match to chrR before performing variant calling; and (2) the effect of mapping either to chrR alone or the whole genome with chrR. We tested all four different combinations of these tasks and found that incorporating MUMmer improves both runtime efficiency and variant calling performance (F1 score) regardless of whether the analysis begins from raw FASTQ files (Fig. 5A) or BAM files (Fig. 5B). In addition, we observed that when the MUMmer step is included, the choice between mapping to just chrR or to the whole reference genome has minimal impact on performance. However, omitting MUMmer, led to an increase of 15 false positives, all mapping to the 18S rRNA gene. These false positives likely originate from pseudogene-derived reads, as 18S pseudogenes are known to be present outside the acrocentric chromosomes (3, 13). This highlights the critical role of the MUMmer-based read retrieval step, which not only accelerates the pipeline runtime but also enhances performance by excluding reads originating from rDNA pseudogenes.

**Figure 5.**
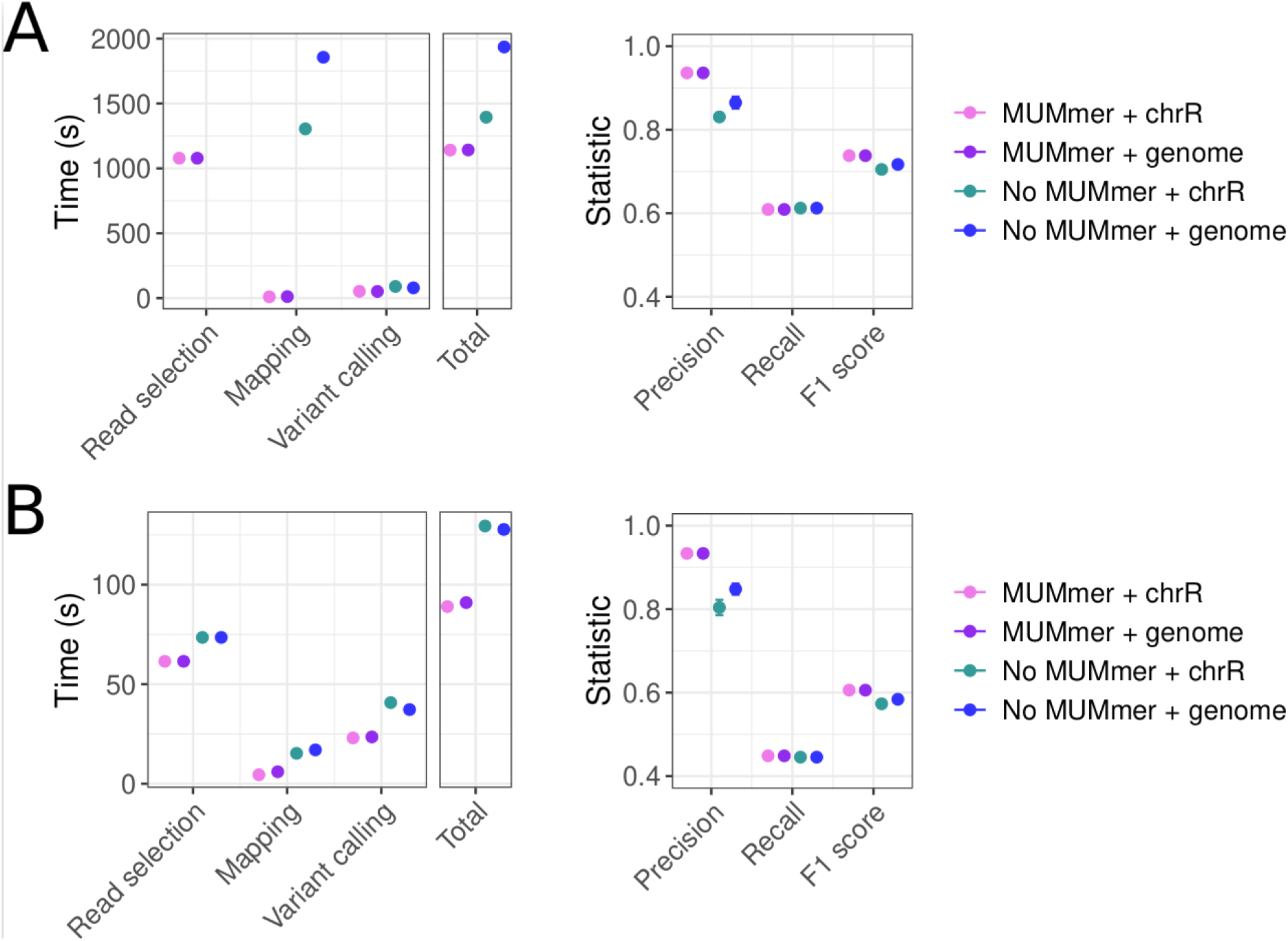
Performance of rDNAcaller on experimental data. Average time and accuracy of *rDNAcaller* on 4 replicates from the CHM13 cell line when the input data is either **(A)** a FASTQ file or **(B)** a bam file already mapped to hg38. Error bars representing the standard error are included. The x-axis refers to the different preprocessing steps of *rDNAcaller*. “Read selection” is the step of retrieving candidate rDNA reads, which in (A) only represents using MUMmer or not, but in (B) it represents retrieving the candidate rDNA reads from the bam file plus the choice of including MUMmer or not. “Mapping” refers to the mapping with bwa to either a reference with only chromosome R (chrR), or to the optimized genome for rDNA containing only one rDNA copy (genome). “Variant calling” refers to the calling with *HaplotypeCaller* from GATK with ploidy 20. Precision, recall and F1 scores are computed considering only SNPs.

### *rDNAcaller* finds thousands of novel rDNA variants across human populations

A previous work found that rDNA variants were positioned nonrandomly in samples from the 1000 Genomes Project (24), with 28S rRNA harbouring more variants than the 18S and the 5.8S rRNAs (6). Since *rDNAcaller* has higher sensitivity than their method (JG - 2, Fig. 2), we expect to find more variants in the same data. Our analysis identified a catalog of 6,019 variant positions, comprising 8,876 variant alleles, ~25% of which are indels (Supplementary Table, Fig. 6A). These variants show significant overlap with those previously detected using the same dataset (6) (OR=5.72, p-value 2.2e-16, Fig. 6B). However, as expected, we retrieved 5,607 additional variant positions that were not reported in the previous study, representing an increase of one order of magnitude. Nearly half of our detected variant alleles (~48%) are individual-specific, whereas 2,124 are shared across five or more individuals.

**Figure 6.**
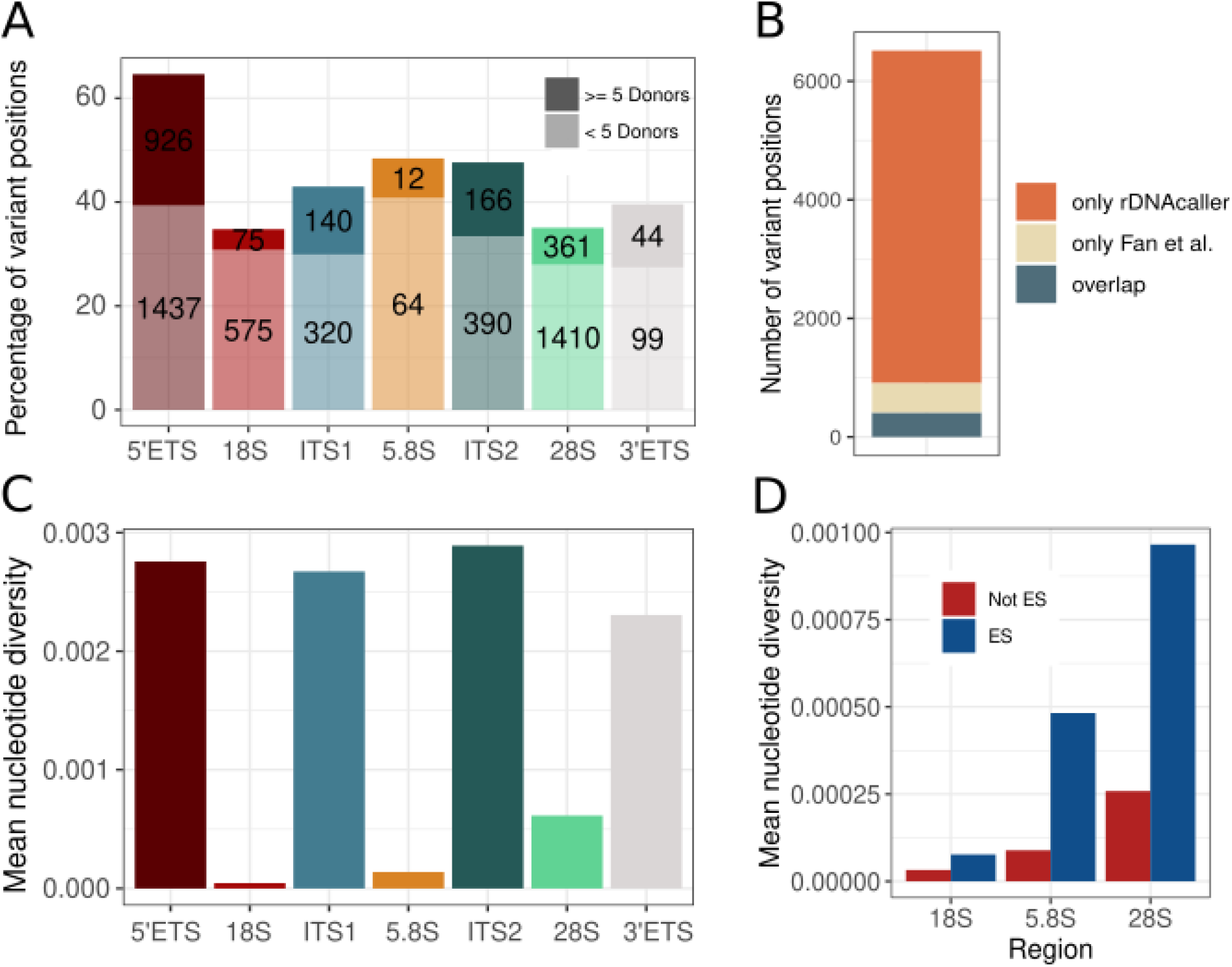
rDNA variants in the 1000 Genomes Project. **A**. Number of rDNA variants stratified by region and by whether they are present in more than 5 donors or not. **B**. Overlap between the variants found with rDNAcaller and (6) **C**. Mean nucleotide diversity stratified by region. **D**. Mean nucleotide diversity stratified by region and location in expansion segments (ES)

We then explored the distribution of the rDNA variants across the different regions of the locus. The 5’ external transcribed spacer (5’ETS) has the highest proportion of variant positions, while the 18S and the 28S rRNAs have the lowest (Fig. 6A). We also computed the average nucleotide diversity score, which accounts for both the proportion of variant positions and their allele frequency. As expected, we found that the three rRNA-encoding regions (28S, 18S and 5.8S) had the lowest nucleotide diversity values, consistent with these regions being more evolutionarily conserved than the spacer regions (Fig. 6C).

Expansion segments are structurally flexible regions of rRNAs that extend beyond the ribosomal core in three-dimensional space. These regions are less conserved across species, and despite interacting with ribosome-associated proteins, their functional roles still remain unclear (27). We found that the nucleotide diversity scores are significantly higher in expansion segments, consistent with previous studies (8), suggesting that these regions experience weaker evolutionary constraints compared to the core ribosomal regions (Fig. 6D).

### African individuals have more rDNA variants

We then wanted to characterize rDNA genetic diversity between human populations. African individuals showed the highest number of rDNA variants among all populations (Fig. 7A). Importantly, African individuals also show a higher rDNA copy number (CN) (Fig. 7B), a variable with a positive correlation to the number of total rDNA variants across individuals (Fig. 7C).

**Figure 7.**
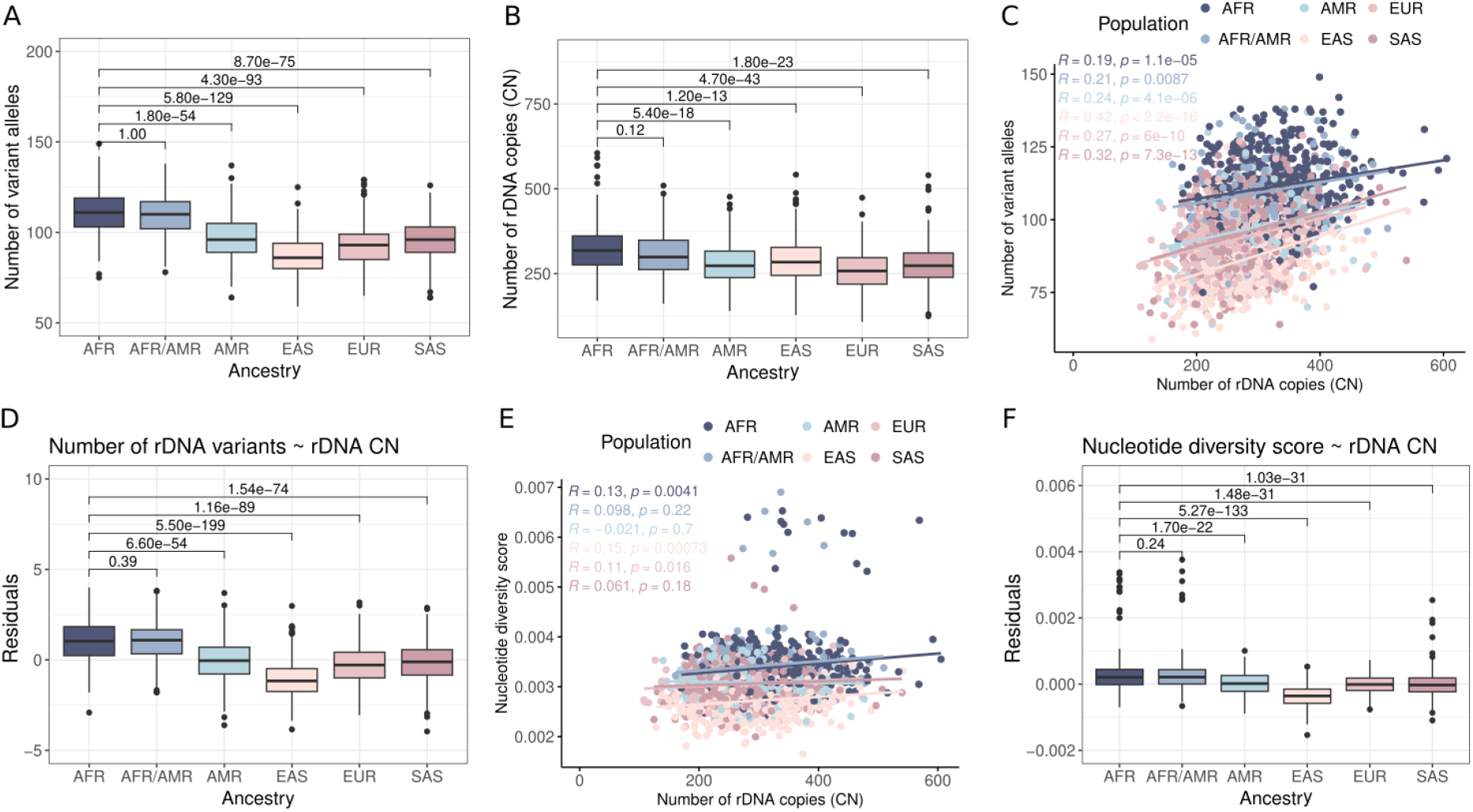
African individuals have a higher number of rDNA variants. **A**. Number of rDNA variant alleles stratified by population (Wilcoxon tests). **B**. rDNA CN stratified by population (Wilcoxon tests). **C**. Correlation between rDNA CN and number of rDNA variant alleles, colored by population. The p-value is obtained from a Pearson correlation. **D**. Residuals obtained from the following linear model: Number of rDNA variants ~ rDNA CN. The p-values are obtained from the covariate Population of the following model: Number of rDNA variants ~ rDNA CN + Population (See Methods). **E**. Correlation between rDNA CN and nucleotide diversity score, colored by population. The p-value is obtained from a Pearson correlation. **F**. Residuals obtained from the following linear model: Nucleotide diversity score ~ rDNA CN. The p-values are obtained from the covariate Population of the following model: Nucleotide diversity score ~ rDNA CN + Population (See Methods). **AFR** = ESN, GWD, LWK, MSL, YRI, ACB, ASW; **AFR/AMR** = ACB, ASW; **AMR** = CLM, MXL, PEL, PUR; **EAS** = CDX, CHB, CHS, JPT, KHV; **EUR** = CEU, GBR, FIN, IBS, TSI; and **SAS** = BEB, GIH, ITU, PJL, STU. All p-values are adjusted for multiple testing using the Bonferroni test correction.

We wondered whether the higher number of variants in Africans is only due to higher CN, or if this variability is still higher when accounting for CN. To check this, we run a linear model adjusting for CN (See Methods), and we see that CN is significantly associated with the number of rDNA variants (p-value=6.35e-46), and African individuals still show a significantly higher number of rDNA variants (Fig. 7D). To further confirm this, we explored the association between population and nucleotide diversity score, a variable less associated with CN (Fig. 7E). In this case, there is also a significant difference between populations, with African individuals showing the highest nucleotide diversity scores. These results show that the higher number of variants in African individuals is not only driven by the higher CN, but African individuals also show greater underlying sequence diversity at rDNA, consistent with their higher effective population sizes and recent evolutionary history (24, 28).

## Discussion

We developed *rDNAcaller*, a fast and highly accurate pipeline for detecting rDNA variants in both simulated and experimental datasets. Unlike previous methods, *rDNAcaller* avoids calling pseudogenic variants, and outperforms them in both accuracy and computational efficiency.

Applying *rDNAcaller* on the 1000 Genomes Project dataset, we successfully replicated previously-identified rDNA variants while also expanding the variant catalog by an order of magnitude. This increase enhances our understanding of human genetic variation within rDNA regions, providing a more comprehensive and accurate representation of these loci. Additionally, we reveal a correlation between higher rDNA CN and increased number of rDNA variants, with African individuals showing the highest values for both. Importantly, after controlling for CN, African individuals still exhibit more rDNA variants and higher nucleotide diversity scores than other populations, consistent with the patterns observed in the rest of the genome, where African populations show greater sequence diversity (24, 28). The high similarity among rDNA copies has been attributed to concerted evolution (29), where multiple copies of a repeat family are homogenized over time to maintain high sequence similarity within a species. However, the persistence of rDNA variants indicates an incomplete concerted evolution (11). Our findings suggest that rDNA variants escaping homogenization reflect the same population-level diversity seen across the genome, with African populations showing the greatest variation. Importantly, a previous study (7) reporting fewer rDNA variants did not detect this trend, highlighting the value of the increased sensitivity of *rDNAcaller*.

We anticipate *rDNAcaller* to render useful in many biological contexts. In genome-wide association studies (GWAS), rDNA variants have traditionally been excluded (2). However, a recent study focused on a very small subset of rDNA variants has already shown important associations with body size (9). Our pipeline will greatly increase the number of rDNA variants to be tested for associations with human traits in large biobanks. Additionally, rDNA copy number is extremely affected by cancer (30), potentially giving rise to novel rDNA variants or altering the frequency of existing variants. By integrating *rDNAcaller* into cancer genomics, we might uncover new insights into the role of rDNA variation in tumorigenesis. Beyond cancer, our pipeline holds promise for studying a plethora of diseases caused by ribosomal dysfunction, named ribosomopathies (31). Some of these conditions may originate from specific rDNA variants, and *rDNAcaller* provides a powerful tool for identifying such pathogenic variants. Notably, some of these variants may arise somatically, since rDNA might be a hotspot for somatic mosaicism, and rDNAcaller could be instrumental in identifying such variation across tissues and disease states. Finally, our pipeline could be adapted for use in other species, such as mice, and other repetitive genomic elements with architectures similar to rDNA, such as SINEs and LINEs, which also show intra- and inter-individual variation.

Overall, *rDNAcaller* represents a significant advancement in the study of human genetic diversity. By enabling more precise analysis of rDNA variants, this pipeline has the potential to enhance our understanding of genetic variation in ribosomal DNA and its implications for health and disease, ultimately bridging the gap between rDNA diversity and phenotypic traits in biomedical research.

## Acknowledgements

J.M.R. was supported by a predoctoral fellowship from “la Caixa” Foundation (ID 100010434) with code LCF/BQ/DR22/11950022. M.M. was supported by a grant PID2019-107937GA-I00 funded by MCIN/AEI/10.13039/501100011033 and a grant RYC-2017-22249 funded by MCIN/AEI/10.13039/501100011033 and by “ESF Investing in your future”.

We thank Pau Clavell-Revelles for insightful discussions on population genetics.

## Author contributions

J.M.R. led the analysis. R.G., M.V., A.J. and A.S., contributed to the simulations. A.S. and P.P. contributed to the CHM13-T2T analysis and Figure 4. W.O. analyzed the 1KGP data and contributed to Figures 6 and 7. A.J. contributed to Figure 1 and 2. M.M. conceived and supervised the work. J.M.R., W.O. and MM wrote the manuscript.

## Data and code availability

The *rDNAcaller* is available at: https://github.com/Mele-Lab/rDNAcaller, and the simulator is available at: https://github.com/Rbbt-Workflows/SyntheticRDNA.

